# The environmental stress response regulates ribosome content in cell cycle-arrested *S. cerevisiae*

**DOI:** 10.1101/2021.05.14.444167

**Authors:** Allegra Terhorst, Arzu Sandikci, Gabriel E. Neurohr, Charles A. Whittaker, Tamás Szórádi, Liam J. Holt, Angelika Amon

## Abstract

Temperature sensitive cell division cycle (*cdc-ts*) cells are unable to progress through the cell cycle at the restrictive temperature due to mutations in genes essential to cell cycle progress. Cells harboring *cdc-ts* mutations increase in cell volume upon arrest but eventually stop growing. We found that this attenuation in growth was due to selective downregulation of ribosome concentration. We saw similar ribosome downregulation in cells arrested in the cell cycle through alpha factor addition, rapamycin addition, and entrance into stationary phase. In all cell cycle arrests studied, cells activated the Environmental Stress Response (ESR), a key transcriptional response to many stressors in *S. cerevisiae*. When we combined cell cycle arrest with hyperactivation of the Ras/PKA pathway, ESR activation was prevented, cells were unable to downregulate their ribosomes, and cell viability was decreased. Our work uncovers a key role for the environmental stress response in coupling cell cycle progression to biomass accumulation.

## Introduction

For organisms to maintain both their fitness as well as their organismal and cellular homeostasis, biomass accumulation and cell division must be tightly coordinated (1, 2). Throughout the G1 phase, the presence of growth signals, amino acids, and glucose stimulates the rate of macromolecule biosynthesis, increasing the cell’s volume. Entry into the cell cycle relies on sufficient biosynthetic capacity to reach a cell’s “critical size,” or minimum size threshold dependent on nutrient availability (3, 4). While cell size control mechanisms seem to prevent cells from entering cell division while too small, there are also issues associated with cells being too large, such as decreased surface area to volume and DNA to cytoplasm ratios. Because alterations of these key cellular parameters can detrimentally affect cell viability, it is likely that cells have evolved to maintain homeostatic size and biomass production (3, 5, 6, 7).

*Saccharomyces cerevisiae*, or budding yeast, has proven to be critical in the study of the relationship between cell growth and progression through the cell cycle. In order to understand this relationship, cell growth and division can be uncoupled using temperature-sensitive cell division cycle (*cdc-ts*) mutants, which have mutations in genes required for cell cycle progression. At the restrictive temperature, the *cdc-ts* mutants are unable to progress through the cell cycle but continue to accumulate biomass and thus increase in size, in some mutant strains up to 16 times the size of a wild-type cell (8, 9). Previous work from our lab determined that, at the restrictive temperature, the size of many *cdc-ts* mutants eventually plateaus (9). In one of these *cdc-ts* mutants, attenuation of cell growth in oversized cells correlated with an unusual dilution of the cytoplasm, suggesting that reduced overall biomass production might cause attenuation of growth during prolonged cell cycle arrests (6).

The ability or inability to produce ribosomes has been shown to cause dramatic changes in cell size (4). A significant portion of a cells’ energy is involved in accumulating biomass through protein synthesis by ribosomes. Ribosomes, which are composed of ribosomal proteins and rRNA, translate mRNA into proteins. The biogenesis of ribosomes is highly regulated within the cell, to prevent cells from unnecessarily expending energy (10). When energy is abundant, cells rapidly make ribosomes, and the ribosomal fraction of the proteome correlates with growth rate (11). We hypothesized that cell cycle arrested cells would shift energy expenditure from growth to maintenance of viability. Using different *cdc-ts* mutants as a model system, we here describe our studies determining how ribosome content regulates cell size in arrested cells.

In this report, we determined how cells regulate growth and biomass production during prolonged cell cycle arrests in budding yeast. Using ribosome purification and Tandem Mass Tag (TMT) Proteomics, we found that ribosomes were specifically downregulated during cell cycle arrests in *cdc-ts* mutants. We saw a similar ability to attenuate size through downregulation of ribosome biogenesis in physiological cell cycle arrests. All investigated cell cycle arrests led to activation of the Environmental Stress Response (ESR), a transcriptional response to stress in *S. cerevisiae* that decreases translational capacity (12, 13). Preventing ESR activation through hyperactivation of the Ras/PKA pathway interfered with downregulation of ribosome content and reduced cell survival during the cell cycle arrest. We conclude that activation of the ESR, potentially through attenuation of ribosome biogenesis, is required for survival of cell-cycle arrested cells.

## Results

### Cell volume is regulated in *cdc-ts* arrested cells

In order to determine how cells attenuate their biomass accumulation during cell cycle arrests, we studied three independent *cdc-ts* mutants: *cdc28-13, cdc20-1*, and *cdc15-2*. We decided to examine these three separate *cdc-ts* mutants because they arrest in three distinct phases of the cell cycle and grow to different maximum cell volumes. This allowed us to distinguish between global, size-specific, and cell cycle phase-specific observations. Cdc28 is the main cyclin-dependent kinase that coordinates the yeast cell cycle, and *cdc28-13* cells arrest in G1 (9, 14). Previous work from our lab has shown that in *cdc28-13* cells, the mean cell volume growth is driven by growth of the whole cell, including the cytoplasm, rather than solely vacuole enlargement (6). Cdc20 is the activator subunit of the anaphase promoting complex that drives the metaphase-anaphase transition, therefore *cdc20-1* mutants arrest in metaphase at the restrictive temperature (9, 15). Cdc15 is a critical kinase required for mitotic exit, and *cdc15-2* mutants at the restrictive temperature arrest in late anaphase (9, 16).

We measured the mean cell volume of all three *cdc-ts* strains was measured using a Coulter Counter to confirm the previous findings of Goranov et al. (2009; 9). In agreement with previous cell size data (9), the mean cell volume of all three *cdc-ts* strains increased over 9 hours and eventually plateaued (*Figure 1A*). As a control, a *WT* strain (*Figure 1A*, gray) was grown concurrently and was kept in log phase, termed “cycling,” to avoid confounding factors caused by growth into stationary phase, which is a response to glucose depletion and known to cause many physiological changes including changes in cell size (17, 18). The cell volume of *cdc28-13* mutant cells grew exponentially for the first two hours of arrest, grew linearly until 6 hours, and eventually plateaued to the largest of the strains, at 800 fL (*Figure 1A*, pink). The mean cell volume of *cdc20-1* cells initially was the largest of the strains and appeared to have two distinct plateaus, one at 300 fL after being arrested for five hours and the other at 450 fL after 9 hours of cell cycle arrest (*Figure 1A*, purple). Cell harboring the *cdc15-2* mutation had linear growth and plateaued to approximately 500 fL (*Figure 1A*, blue). We note that both *cdc20-1* and *cdc15-2* strains have final sizes that differ from those previously reported in Goranov et al. (2009; 9), likely due to the slightly differing methods used to measure cell size.

**Figure 1.**
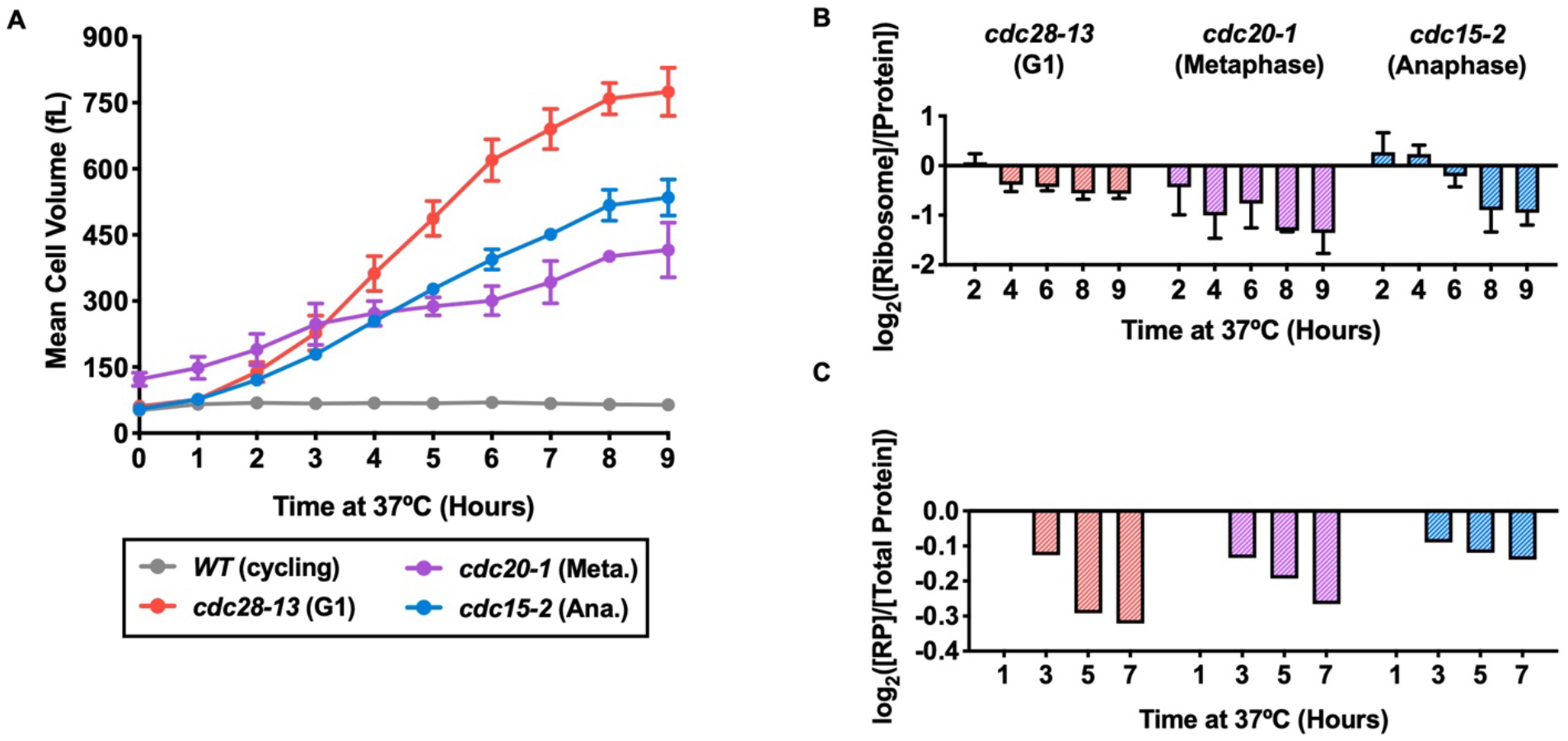
Ribosome concentrations decrease during prolonged cell cycle arrest. *WT* (gray, A2587), *cdc28-13* (red, A39000), *cdc20-1* (purple, A937), and *cdc15-2* (blue, A2596) cells were grown to log phase in YEPD at 25°C and then shifted to 37°C for 9 hours. *WT* cultures were kept in log phase (cycling) at OD_600nm_ 0.2-0.8, by diluting with pre-warmed (37°C) YEPD. ***(A)*** Mean cell volume (fL) was measured. Error bars represent standard deviation from the mean of experimental replicates. ***(B)*** Protein and ribosome concentrations were quantified using a quantitative ribosome purification method, described in Terhorst et al. (2020; 17). Samples of equal volume were lysed with a French Press, and [Protein] was measured with a Bradford Assay. Intact ribosomes were purified with sucrose cushion centrifugation, and [Ribosome] was measured by rRNA absorbance at A260nm on a Nanodrop. [Ribosome]/[Protein] were determined. Values were normalized to those of the *WT* cycling samples at each time point and subsequently log_2_ transformed. Error bars represent standard deviation from the mean of experimental replicates. ***(C)*** TMT Proteomics was performed on the *cdc-ts* strains. *cdc28-13* cells were synchronized with alpha factor prior to arrest at 37°C. The fraction of ribosomal proteins (RP) in total protein extracts ([RP]/[Total Protein]) was determined. Values were normalized to the 1-hour time point in each experiment and subsequently log_2_ transformed.

### Protein and ribosomes are downregulated in size-attenuated *cdc-ts* arrested cells, leading to increased cytoplasmic diffusion

Having confirmed that cell growth is attenuated during various *cdc-ts* mutant cell cycle arrests, independent of cell cycle stage in which the cells are arrested, we hypothesized that this global size control was regulated by cellular ribosome content because it is known that ribosome biogenesis is rate-limiting for growth in yeast (11). To test this hypothesis, we measured the protein and ribosome content of the *cdc-ts* mutant strains during 9-hour cell cycle arrests to determine protein and ribosome concentrations as cell growth plateaued. Cells were lysed, and protein concentration ([Protein]) of the lysates was measured with a Bradford Assay. We next measured the rRNA concentration of intact ribosomes ([Ribosome]) purified with sucrose-cushion centrifugation. To measure the ribosomal fraction of the proteome, we next calculated [Ribosome]/[Protein] from the previous measurements. Because the arrest of *cdc-ts* strains requires a temperature shift from 25°C to 37°C, we began protein and ribosome measurements after 2 hours at 37°C to avoid confounding effects of heat shock (19, 20). Concurrent to the arrests of the *cdc-ts* mutants, we also grew a *WT* cycling culture and normalized each *cdc-ts* time point to those of the *WT* cycling culture. Because ribosome biosynthesis rate is known to be a major determinant of cell size, we expected to see a plateau or decrease in the ribosome content of the *cdc-ts* cells compared to *WT* cycling cells (4, 11). Consistent with this prediction, [Ribosome]/[Protein] decreased dramatically over the 9-hour cell cycle arrests in all three *cdc-ts* strains, (*Figure 1B*). We confirmed these findings by measuring the fraction of the proteome consisting of ribosomal proteins ([RP]/[Total Protein]) using TMT proteomics of whole cell lysates. Again, we saw a decrease in [RP]/[Total Protein] in all three *cdc-ts* arrests (*Figure 1C*). The data from these two separate methods, taken together, suggest a specific downregulation of ribosomes during *cdc-ts* cell cycle arrests, which is independent of the specific cell cycle arrest, while cells continue to make other protein. We hypothesize that the resulting decreased translational capacity leads to attenuation of cell volume growth after prolonged cell cycle arrest.

Cytoplasmic ribosome concentration not only affects translational capacity but also contributes to molecular crowding and thereby indirectly influences important processes, such as phase separation and cytoplasmic diffusion (21). To determine whether cytoplasmic crowding was affected in *cdc-ts* cell cycle arrests, we used genetically encoded multimeric nanoparticles (GEMs) to probe the mesoscale diffusivity of the cytoplasm as described in Delarue et al. (2018, 21). Upon mTORC1 inactivation using rapamycin, Delarue et al. (2018, 21) reported a significant increase in the cytoplasmic effective diffusion coefficient of the GEMs, which was attributed to the presence of fewer ribosomes in the cytoplasm (21). The median effective diffusion coefficients of GEMs in *cdc-ts* and *WT* cycling cells were measured at different time points over 6 hours, at which point ribosome and protein content are downregulated in *cdc-ts* mutants. The median effective diffusion coefficients of the *cdc-ts* and *WT* cycling cells increased considerably between 0 hours and 2 hours, during which they were shifted to 37°C. We attribute this increase to heat shock, which has been previously shown to increase cytoplasmic diffusion (11). After the initial heat shock, the median diffusion coefficients of all three *cdc-ts* mutants remained above the median diffusion coefficient of the *WT* cycling cells, which decreased gradually after heat shock (*Figure S1F*). As expected, *cdc20-1* cells had the highest median diffusion coefficient initially and throughout the experiment, in agreement with our observation that *cdc20-1* cells are already downregulating ribosomes at the permissive temperature. We conclude that ribosome downregulation causes decreased macromolecular crowding of the cytoplasm in *cdc-ts* cells arrested in different cell cycle phases.

### G1 arrests decrease ribosome content to attenuate cell growth, independent of the method used for the arrest

Whether growth attenuation caused by ribosome downregulation is a global phenomenon remained unknown, so we next studied changes in cell size and protein and ribosome content in naturally occurring cell cycle arrests. We investigated the growth of cells arrested with three different methods: alpha factor pheromone addition, TORC1 inhibition through rapamycin addition, and entry into stationary phase due to glucose depletion. Alpha factor (αf) is a pheromone secreted by mating-type alpha (*MATα*) cells that arrests mating-type a (*MATa*) in G1 before mating begins through a well-studied MAPK pathway that previous work from our lab has shown to inactivate TORC1 (22, 23). Rapamycin addition is also known to arrest cells in G1 by directly inhibiting TORC1, mimicking the effect of amino acid starvation (4, 24, 25). Depletion of glucose causes inhibition of both TORC1 and a related nutrient-sensing pathway, the Ras/PKA pathway. Cells started to enter stationary phase upon growth to saturation at OD_600nm_ of approximately 3.4 in our spectrophotometer. At this point, cells become starved for glucose, causing arrest in G1 (4, 24). These three G1 arrests function through different mechanisms and therefore, were used to determine whether ribosome downregulation only occurs in *cdc-ts* arrested cells or is independent of the method used to arrest cells.

We first measured mean cell volume throughout each cell cycle arrest. *bar1Δ* cells were used in the alpha factor experiments. These cells lack the protease that degrades alpha factor, thus ensuring that alpha factor was not degraded and that cells did not escape their cell cycle arrests. *bar1Δ* cells arrested with alpha factor (*Figure 2A*, +αf, orange) for 9 hours reached a maximum mean cell volume of over 200 fL; in comparison to *bar1Δ* cells grown in the absence of alpha factor (*Figure 2A*, −*α*f, yellow), cells arrested with alpha factor grew to be four times larger (26). After 8 hours arrest, alpha factor arrested cells began to slow in cell growth at a cell volume around 200 fL, which was a smaller maximum mean cell volume than in the *cdc28-13* G1 arrested cells, suggesting the presence of size attenuation during the arrest (*Figure 2A*). *WT* haploid cells grown into stationary phase (*Figure 2B*, green squares) reached an OD_600nm_ of 8.5 by 10 hours and of 20.0 by 24 hours. The mean cell volume of these cells decreased as cells became starved for glucose and entered stationary phase, suggesting a potentially interesting relationship between glucose starvation and size attenuation (*Figure 2B*). In rapamycin experiments, *WT* haploid cells were arrested with rapamycin for 5 hours (*Figure 2C*, green circles). Although TORC1 inhibition has been shown to decrease cell size in mammalian cells (27), there was a subtle increase in mean cell volume throughout the rapamycin arrest in the W303 *S. cerevisiae* cells, reaching a final volume of approximately 130 fL, twice that of an untreated cycling *WT* haploid cell (*Figure 2C*). The increase in mean cell volume upon rapamycin addition could be explained by growth of the vacuole, which is known to occur in *S. cerevisiae* after rapamycin addition, and the concentration of the cytoplasm specifically may be changing (28). Although the mean cell volume of rapamycin arrested cells increased, size attenuation may have prevented these cells from growing even larger. With the exception of the rapamycin experiments, cell size decreased in the naturally occurring cell cycle arrests, suggesting that growth attenuation during cell cycle arrests does not only happen in *cdc-ts* mutants and is independent of method used to arrest the cells.

**Figure 2.**
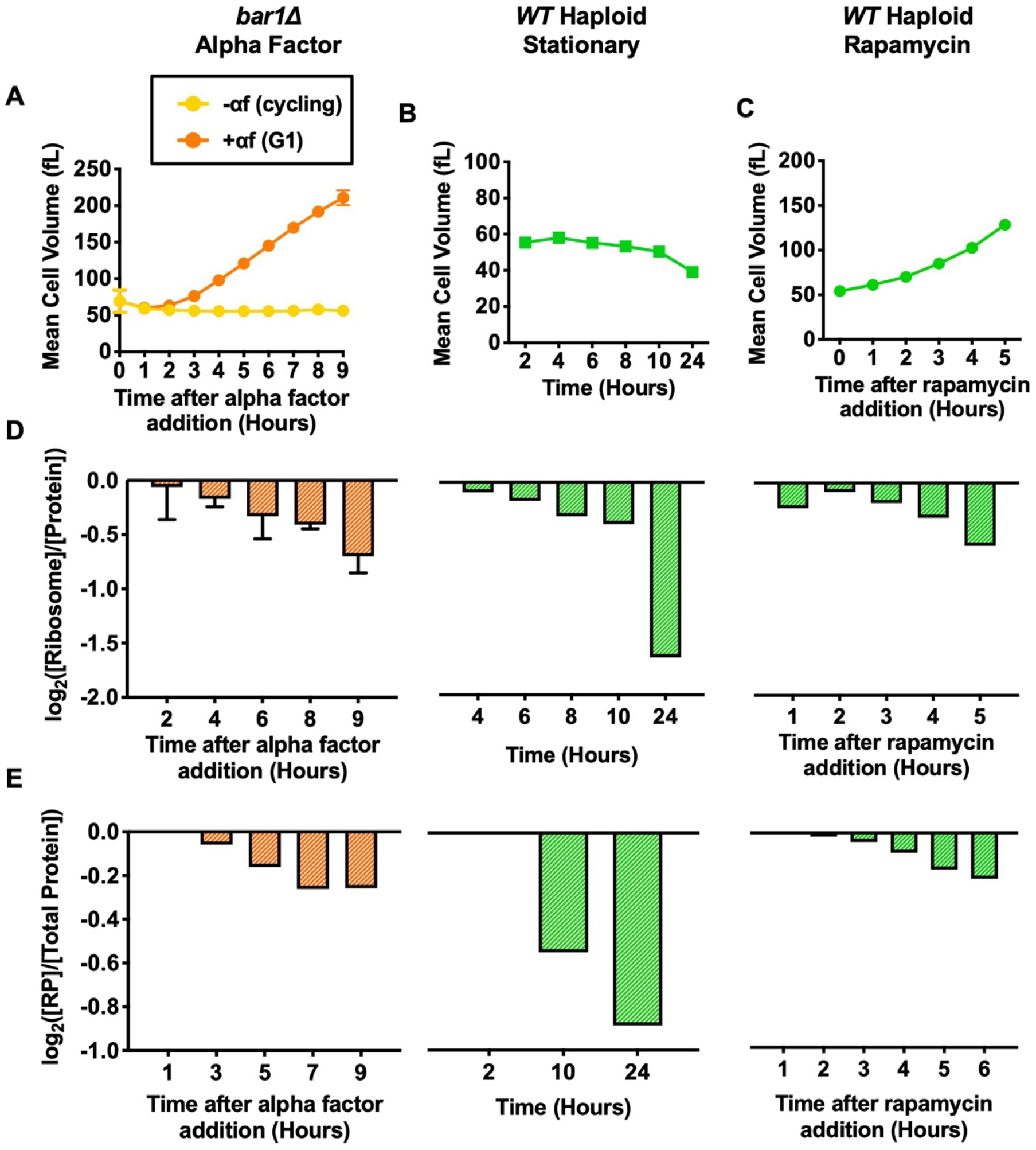
Protein and ribosome quantification of cells arrested in G1 with various methods. For alpha factor experiments, *bar1Δ* (A2589) cells were grown to log phase in YEPD at 30°C. Cells were then divided into two cultures and grown for 9 hours at 30°C. 5 μg/mL alpha factor was added to one culture (+αf, orange) while the equivalent volume of DMSO was added to the other (−αf, yellow). 2 μg/mL of alpha factor (+αf, orange) or the equivalent volume of DMSO (−αf, yellow) was re-added every 2 hours. For stationary phase experiments, *WT* haploid (green, A2587) cells were grown in YEPD for 24 hours at 30°C. For rapamycin experiments, *WT* haploid (green, A2587) cells were grown to log phase in YEPD at 30°C. 5 nM rapamycin was added, and cells were grown for 5 hours at 30°C. ***(A-C)*** Mean cell volume (fL) was measured for ***(A)*** *bar1Δ* cells with (+αf, orange) and without (−αf, yellow), ***(B)*** *WT* haploid cells grown into stationary phase (green squares), and ***(C)*** *WT* haploid cells with added rapamycin (green circles). Error bars represent standard deviation from the mean of experimental replicates. ***(D)*** Protein and ribosome concentrations were quantified using the method described in Terhorst et al. (2020; 17). [Ribosome]/[Protein] was determined. Values were normalized to those of the cycling samples in each experiment (−αf in alpha factor experiments, 2-hour time point in stationary phase experiments, and 0-hour time point in rapamycin experiments) and subsequently log_2_ transformed. Error bars represent standard deviation from the mean of experimental replicates. ***(E)*** TMT Proteomics was performed for each naturally occurring cell cycle arrest. The fraction of ribosomal proteins (RP) in total protein extracts ([RP]/[Total Protein]) was determined. Values were normalized to the 1-hour time point in each experiment and subsequently log_2_ transformed.

To investigate how cell growth was attenuated in natural cell cycle arrests, we determined whether ribosome concentration was decreasing using the method from Terhorst et al. (2020, 17). Indeed, each arrest saw a dramatic decrease in [Ribosome]/[Protein], suggesting that ribosomes are selectively downregulated during cell cycle arrests independent of the method used to arrest cells (*Figure 2D*). This decrease in [Ribosome]/[Protein] was confirmed with TMT Proteomics (*Figure 2E*). As in the *cdc-ts* measurements, the percent change was slightly different between the two methods due to differences in normalization and [Ribosome] measuring concentration of purified assembled ribosome, while [RP] measured the concentration of all ribosomal proteins in whole cell lysates. Still, both experiments suggest that ribosome downregulation occurs in cell cycle arrests independent of method used to arrest the cells.

To ensure the accuracy of our protein and ribosome measurements, the protein and ribosome contents of *WT* haploid (closed green; *Table S1*, A2587) and *WT* diploid (open dark green; *Table S1*, A33728) strains were compared, specifically in the stationary phase and rapamycin experiments. The mean cell volume of the *WT* diploid cells was approximately triple that of the *WT* haploid cells throughout both experiments, particularly when cells reach stationary phase (*Figures S2A-B*). In the stationary phase experiments, the mean cell volume *WT* haploid cells (*Figure S2A*, closed green squares) decreased gradually as cells entered stationary phase, but *WT* diploid cells (*Figure S2A*, open dark green squares) had a sharp decrease in mean cell volume after 8 hours growth, at OD_600nm_ of 3.8. When cells were arrested with rapamycin, the cell volume of both *WT* haploid (*Figure S2B*, closed green circles) and *WT* diploid (*Figure S2B*, open dark green circles) increased exponentially throughout the G1 arrest. Each value was normalized to the 1-hour time point of the *WT* haploid strains to better represent how the two strains differ. In the two experiments, [Ribosome]/[Protein] was the same for *WT* haploid and *WT* diploid strains at each time point, suggesting that the two have the same ribosomal fraction of the proteome (*Figure S2C*). Taken together, the relative measurements [Ribosome]/[Protein] in *WT* haploid and *WT* diploid cells support the accuracy of our measurements.

### Cell cycle arrested cells activate the Environmental Stress Response

We next wanted to understand how the cell is able to downregulate ribosomes in response to cell cycle arrests. Our previous work showed that slow growing cells activate an Environmental Stress Response (ESR) to a degree that is correlated to their growth rate (17, 29), so we collected RNASeq data to determine if the ESR is activated in our arrested cells. The ESR is a transcriptional response to a variety of stresses, such as oxidative and reductive stresses, heat shock, hyperosmotic shock, proteotoxic stress, and nutrient limitation, and was originally described by Gasch et al. (2000; 12). There are approximately 300 genes upregulated in the ESR, a gene set termed the “induced ESR” and involved in promotion of cell survival in stressful conditions by increasing autophagy, DNA damage repair, cell wall reinforcement, and protein folding and degradation. Over 600 genes are downregulated in the ESR, forming a gene set known as the “repressed ESR.” The repressed ESR downregulates many genes related to size and biomass accumulation, such as ribosomal biogenesis genes, RNA production and processing genes, and protein synthesis genes (12, 13).

Previous work from our lab has shown a strong correlation between slow growth and induction of the ESR in yeast strains that have gained and/or lost one or more chromosomes, a condition termed aneuploidy (17, 29). In these aneuploid strains, we have also seen that activation of the ESR, specifically the repressed ESR, which contains many ribosomal protein and biogenesis genes, correlates with a decrease in ribosome content (17). The slow growth and cell stress associated with the *cdc-ts* arrests may lead to activation of the ESR and, consequently, downregulation of ribosomal transcription.

To investigate whether the ESR is activated in cell cycle arrests, we performed RNASeq experiments and analyzed the transcriptomes using a single-sample Gene Set Enrichment Analysis (ssGSEA), described in Tarca et al. (2013, 30) and Terhorst et al. (2020, 17). This analysis calculates ssGSEA projection values for each sample to measure the changes in gene expression distribution of the ESR gene set produced by Gasch et al. (2000; 12). The ssGSEA projection values were calculated separately for the induced ESR and the repressed ESR. As a positive control of ESR activation, *WT* cells were grown in the presence of 500 mM NaCl for 40 minutes to cause hyperosmotic shock (GSE146791, accession numbers: GSM4407212, GSM4407213, and GSM4407214; Terhorst et al. (2020, 17)) (17, 31).

In all three *cdc-ts* arrests, the induced ESR (*Figure 3A*) and the repressed ESR (*Figure 3B*) ssGSEA projection values approach or surpass that of the positive control during the 9-hour arrests. During the cell cycle arrest, cells containing the *cdc28-13* mutation increased their induced ESR ssGSEA projection values above the positive control (*Figure 3A*, red), while dramatically decreasing their repressed ESR ssGSEA projection values far below the positive control (*Figure 3B*, red). *cdc20-1* mutant cells slightly exhibit both the induced ESR (*Figure 3A*, purple) and repressed ESR (*Figure 3B*, purple) at the beginning of the experiment, and its ssGSEA projection values near that of the positive control throughout the arrest. The early activation of the ESR at the 2-hour time points of the *cdc20-1* arrests suggest that the ESR may be already activated in *cdc20-1* cells at the permissive temperature. Cells with the *cdc15-2* mutation exhibit the ESR during their arrest as well. The induced ESR ssGSEA of *cdc15-2* cells reached the positive control by 6 hours, while the repressed ESR ssGSEA decreased below the positive control by 8 hours (*Figure 3A-B*, blue). Since it is known that a large portion of the reduced ESR is composed of ribosomal protein and ribosome biogenesis genes, the ESR activation seen in *cdc-ts* mutants is likely responsible for the selective ribosome downregulation in these strains (12).

**Figure 3.**
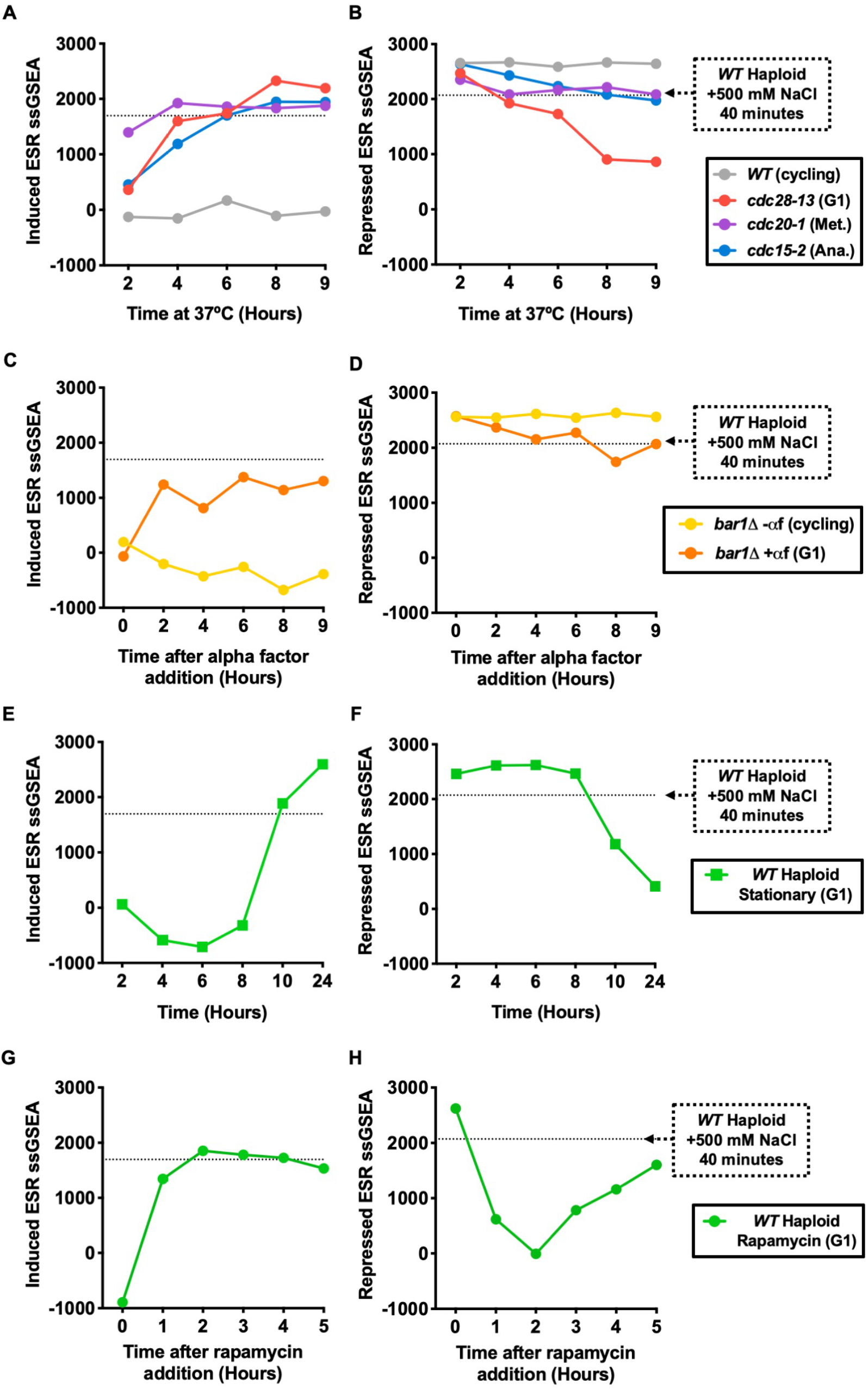
The Environmental Stress Response (ESR) is activated in cell cycle arrested cells. ***(A-B)*** *WT* haploid (gray, A2587), *cdc28-13* (red, A39000), *cdc20-1* (purple, A937), and *cdc15-2* (blue, A2596) cells were grown to log phase in YEPD at 25°C and then shifted to 37°C for 9 hours. *WT* cultures were kept in log phase, termed cycling, at OD_600nm_ 0.2-0.8, by diluting with pre-warmed (37°C) YEPD. RNA-Seq samples were collected, and gene expression data were analyzed by calculating ssGSEA projection values for the **(*A*)** induced ESR and ***(B)*** repressed ESR. The horizontal lines represent the induced ESR and repressed ESR ssGSEA projection values for *WT* cells (A2587) treated with 500 mM NaCl for 40 minutes, a positive control for induction of the ESR. ***(C-D)*** *bar1Δ* (A2589) cells were grown to log phase in YEPD at 30 °C. Cells were then divided into two cultures and grown for 9 hours at 30°C. 5 μg/mL alpha factor was added to one culture (+αf, orange) while the equivalent volume of DMSO was added to the other (−αf, yellow). 2 μg/mL of alpha factor (+αf, orange) or the equivalent volume of DMSO (−αf, yellow) was re-added every 2 hours. RNA-Seq samples were collected, and gene expression data were analyzed by calculating ssGSEA projection values for the **(*C*)** induced ESR and ***(D)*** repressed ESR. The horizontal lines represent the induced ESR and repressed ESR ssGSEA projection values for *WT* cells (A2587) treated with 500 mM NaCl for 40 minutes, a positive control for induction of the ESR. ***(E-F)*** *WT* haploid (A2587) cells were grown in YEPD for 24 hours at 30°C. RNA-Seq samples were collected, and gene expression data were analyzed by calculating ssGSEA projection values for the **(*E*)** induced ESR and ***(F)*** repressed ESR. The horizontal lines represent the induced ESR and repressed ESR ssGSEA projection values for *WT* cells (A2587) treated with 500 mM NaCl for 40 minutes, a positive control for induction of the ESR. ***(G-H)*** *WT* haploid (A2587) cells were grown to log phase in YEPD at 30°C. 5 nM rapamycin was added, and cells were grown for 5 hours at 30°C. RNA-Seq samples were collected, and gene expression data were analyzed by calculating ssGSEA projection values for the (*E*) induced ESR and *(F)* repressed ESR. The horizontal lines represent the induced ESR and repressed ESR ssGSEA projection values for *WT* cells (A2587) treated with 500 mM NaCl for 40 minutes, a positive control for induction of the ESR.

While the ESR has been thoroughly studied after rapamycin addition and growth into stationary phase, we next wanted to confirm whether the ESR occurs in our naturally occurring cell cycle arrests. In the alpha factor experiment, cells grown in the absence of alpha factor did not exhibit either the induced or repressed ESR (*Figure 3C-D*, −αf, yellow), while cells grown in the presence of alpha factor exhibit both the induced and repressed ESR (*Figure 3C-D*, +αf, orange). Previous work from our lab has shown that alpha factor addition causes TORC1 inhibition involving the Fus3 and Kss1 MAPK pathways, raising the possibility that TORC1 inhibition, rather than the cell cycle arrest itself, causes ESR activation in alpha factor-treated cells (23). Cells grown into stationary phase are also known to exhibit the ESR, as glucose starvation causes inhibition of the Ras/PKA pathways (17, 24). As expected, we saw ESR activation in stationary phase cells. This activation was stronger than the positive control by the 10-hour time point and dramatically stronger by the 24-hour time point (*Figure 3E-F*, squares). Rapamycin addition, known to directly inhibit TORC1, caused ESR activation in cells as well (*Figure 3G-H*, circles).

Inhibition of either the TORC1 pathway or the Ras/PKA pathway likely caused the ESR in these naturally occurring cell cycle arrests and lead to the downregulation of their ribosomes.

To further understand the Environmental Stress Response, particularly in our W303 *cdc-ts* mutant strains, we used prototrophic and auxotrophic *cdc28-13* strains to compare the ESR in the presence and absence, respectively, of the ability of a cell to produce a subset of their amino acids. Amino acid starvation through the TORC1 pathway is known to cause activation of the ESR (12). The auxotrophic strain has mutations in genes required for the biosynthesis of leucine, uracil, tryptophan, and histidine, creating exacerbated nutrient starvation conditions, while the prototrophic strain is able to produce these amino acids. Although the two strains were generally the same size when arrested in G1 at the restrictive temperature of 37°C (*Figure S3A*), prototrophic *cdc28-13* cells exhibited the induced ESR (*Figure S3B*) and the repressed ESR (*Figure S3C*) much less than auxotrophic *cdc28-13* cells, particularly after the 4-hour time point. One explanation for the difference in ESR strength is that as the *cdc28-13* cells increase their cell volume, their surface area to cell volume ratio decreases, decreasing the cells’ ability to uptake nutrients, particularly amino acids, and begin to internally starve (28, 32). Auxotrophic *cdc28-13* cells cannot produce their own amino acids and so may be more starved of nutrients than equivalent prototrophic strains. Because the auxotrophic strain exhibits the ESR more robustly than the prototrophic strain, we conclude that as auxotrophic *cdc-ts* cells become too large, the ESR is further activated potentially due to an internal nutrient starvation.

### Cells with a hyperactive Ras/PKA pathway do not exhibit the ESR, leading to a loss of viability

To understand the role of the ESR in *cdc-ts* cell cycle arrests, we wanted to observe the effects of preventing ESR activation during these arrests. Hyperactivation of the Ras/PKA pathway dramatically changes the localization of ESR-dependent transcription factors, suggesting that Ras/PKA hyperactivation suppresses ESR activation (33). In order to hyperactivate the Ras/PKA pathway, we depleted Bcy1, a direct inhibitor of the *S. cerevisiae* PKA orthologues, Tpk1-3, using an auxin-inducible degron (AID) (34). Upon auxin addition, AID-Bcy1 is degraded by the proteasome. Previous work from our lab has further characterized this construct and shown rapid degradation of AID-Bcy1 within 90 minutes of auxin addition (35).

We set out to determine whether the ESR activation in *cdc-ts* cell cycle arrests could be suppressed by hyperactivation of PKA. ESR activation was strongly apparent in *cdc-ts* cells after 4 hours of cell cycle arrest at 37°C (*Figure 3A-B*). Therefore, we aimed to compare the ESR with and without BCY1 depletion at the 4-hour time point.

We concurrently grew *cdc-ts* cells without the *AID-BCY1* construct. A recent publication from our lab shows convincing data suggesting that the ESR is required to recover from the heat shock response (35), and therefore, we waited for 2 hours after heat shock before adding auxin to allow the general effects of heat-shock to resolve and thus prevent heat-shock related cell death. Auxin was re-added every two hours to maintain depletion of Bcy1 throughout the 6-hour arrest, and *WT* and *WT AID-BCY1* cells were kept in log-phase. As before, we compared iESR and rESR values to those of cells experiencing hyperosmotic shock as a positive control.

As before (*Figure 3A-B*, gray), *WT* cycling cells maintained a constant iESR ssGSEA projection value below the positive control and a constant rESR ssGSEA projection value above the positive control throughout the 6-hour arrest, suggesting that ESR activation does not occur in *WT* cycling cells (iESR, *Figure 4A-C*, gray; rESR, *Figure 4D-F*, gray). Cycling *WT AID-BCY1* cells had lower iESR ssGSEA projection values and higher rESR ssGSEA projection values than cycling *WT* (iESR, *Figure 4A-C*, black; rESR, *Figure 4D-F*, black). These data suggest that hyperactivation of the Ras/PKA pathway suppress ESR activation. While we saw activation of both the iESR and rESR in *cdc28-13* cells that reached that of the positive control (iESR, *Figure 4A*, red; rESR, *Figure 4D*, red), *cdc28-13 AID-BCY1* cells had iESR and rESR ssGSEA projection values resembling those of the cycling *WT* cells (iESR, *Figure 4A*, dark red; rESR, *Figure 4D*, dark red). In *cdc20-1* cells arrested for 6 hours, iESR ssGSEA values increased to that of the positive control, and rESR ssGSEA values decreased gradually toward that of the positive control (iESR, *Figure 4B*, purple; rESR, *Figure 4E*, purple). When *cdc20-1 AID-BCY1* cells were arrested, iESR and rESR ssGSEA projection values were most similar to those of the cycling *WT* cells (iESR, *Figure 4B*, dark purple; rESR, *Figure 4E*, dark purple). *cdc15-2* cells had the most subtle activation of the ESR with its iESR and rESR ssGSEA projection values nearing but not reaching those of the positive control by 6 hours of cell cycle arrest (iESR, *Figure 4C*, blue; rESR, *Figure 4F*, blue). As expected, *cdc15-2 AID-BCY1* cells did not exhibit ESR activation, and iESR and rESR ssGSEA projection values were most similar to those of *WT AID-BCY1* (iESR, *Figure 4C*, dark blue; rESR, *Figure 4C*, dark blue). In conclusion, ESR activation was prevented in all cell cycle arrests when Ras/PKA was hyperactivated.

**Figure 4.**
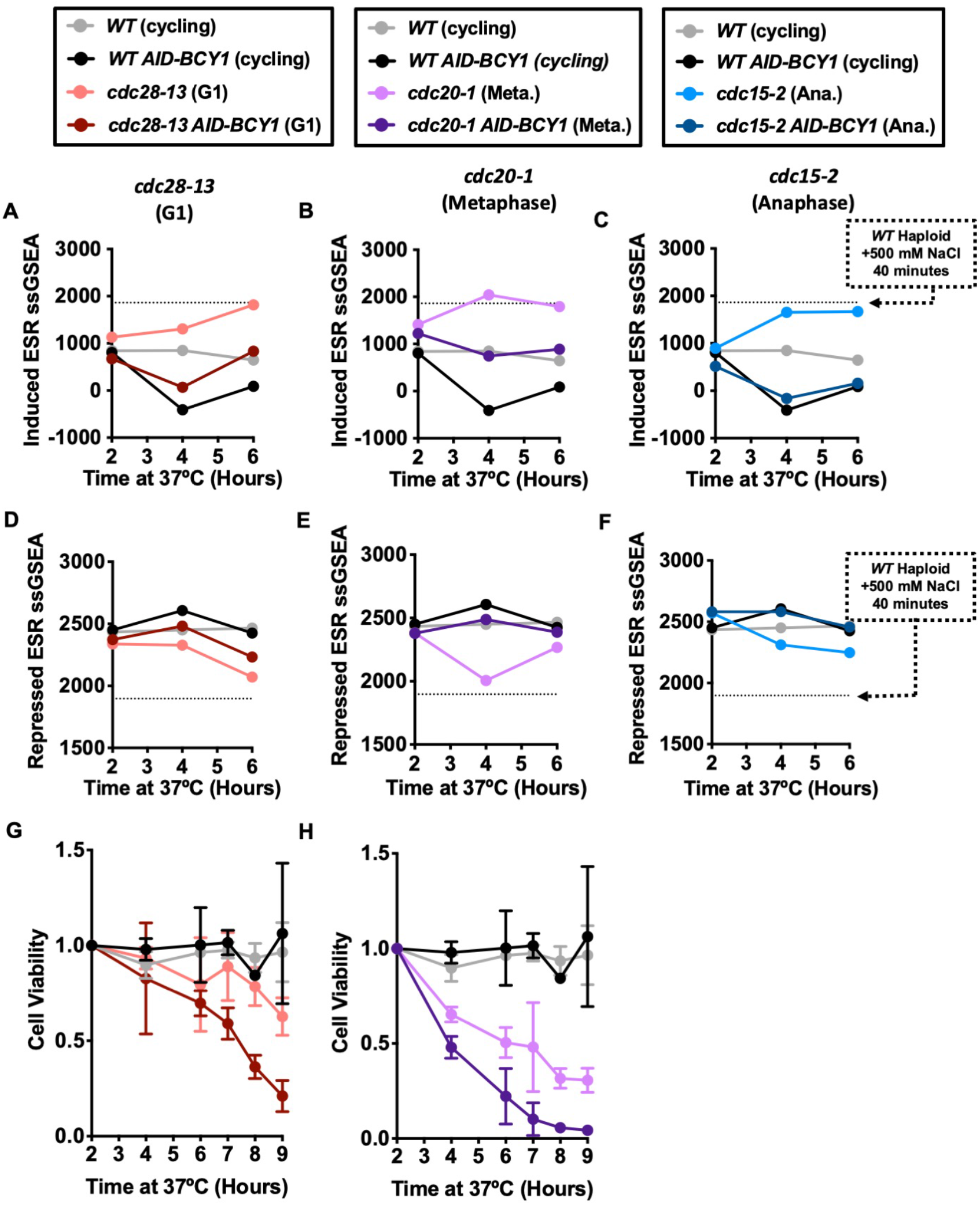
Hyperactivation of the Ras/PKA pathway suppresses the ESR and reduces the viability of cell cycle-arrested cells. *WT* (gray, A2587), *WT AID-BCY1* (black, A40439), *cdc28-13* (red, A39000), *cdc28-13 AID-BCY1* (dark red, A40444), *cdc20-1* (purple, A937), *cdc20-1 AID-BCY1* (dark purple, A40499), *cdc15-2* (blue, A2596), and *cdc15-2 AID-BCY1* (dark blue, A40501) cells were grown to log phase in YEPD (supplemented with 138 μL glacial acetic acid for each 1 L YEPD) at 25°C and then shifted to 37°C for 6 hours. *WT* and *WT AID-BCY1* cultures were kept in log phase, termed cycling, at OD_600nm_ 0.2-0.8, by diluting with pre-warmed (37°C) YEPD (supplemented with 138 μL glacial acetic acid for each 1 L YEPD). 500 μM indole-3-acetic acid was added after 2 hours and 4 hours at 37°C. ***(A-C)*** Induced ESR ssGSEA projection values were calculated from RNA-Seq gene expression data for *WT* (gray), *WT AID-BCY1* (black), and ***(A)*** *cdc28-13* (red) and *cdc28-13 AID-BCY1* (dark red), ***(B)*** *cdc20-1* (purple) and *cdc20-1 AID-BCY1* (dark purple), and ***(C)*** *cdc15-2* (blue) and *cdc15-2 AID-BCY1* (dark blue). ***(D-F)*** Repressed ESR ssGSEA projection values were calculated from RNA-Seq gene expression data for the repressed ESR for *WT* (gray), *WT AID-BCY1* (black) and ***(D)*** *cdc28-13* (red) and *cdc28-13 AID-BCY1* (dark red), ***(E)*** *cdc20-1* (purple) and *cdc20-1 AID-BCY1* (dark purple), and ***(F)*** *cdc15-2* (blue) and *cdc15-2 AID-BCY1* (dark blue). ***(G-H)*** Cell viability was measured for *WT* (gray), *WT AID-BCY1* (black) and *(G) cdc28-13* (red) and *cdc28-13 AID-BCY1* (dark red) and *(H) cdc20-1* (purple) and *cdc20-1 AID-BCY1* (dark purple). Values were normalized to the 0-hour time point of each experiment. Error bars represent standard deviation from the mean of experimental replicates.

We next asked if ESR activation was important for viability during prolonged cell cycle arrest. We grew *WT* and *cdc-ts* cells with and without the *AID-BCY1* construct for 9 hours at 37°C and plated 300 cells on YPD plates at 25°C, the permissive temperature, to determine the viability of cells when they return to the cell cycle. Colonies were allowed to grow for 3 days and were then counted. Cell viability was normalized to the 2-hour timepoint. Unfortunately, the sonication required to accurately determine cell concentration on the Coulter Counter appeared to separate the bud from the mother cell in *cdc15-2* cells once they began to arrest in anaphase. This caused all cells experiencing the *cdc15-2* arrest to die once sonicated, and therefore, *cdc15-2* and *cdc15-2 AID-BCY1* were not included in this analysis. *WT* and *WT AID-BCY1* did not have significantly different cell viability through the 9-hour cell cycle arrest (*Figure 4G-H*, gray and black, respectively). In contrast, *cdc28-13* cells had a cell viability of above 0.5 during the 9-hour arrest, while *cdc28-13 AID-BCY1* cells had fewer cells survive with a cell viability below 0.3 after being arrested for 9 hours (*Figure 4G*, red and dark red, respectively). Similarly, fewer *cdc20-1 AID-BCY1* cells survived throughout the 9-hour arrest than *cdc20-1* cells (*Figure 4H*, dark purple and purple, respectively), but *cdc20-1* cells were less viable than *cdc28-13* cells throughout the arrest. We conclude that the inability to activate the ESR through hyperactivation of the Ras/PKA pathway leads to increased cell death in during cell cycle arrest.

### Hyperactivation of the Ras/PKA pathway prevents size attenuation and ribosome downregulation in *cdc-ts* arrests

The question that remained was whether hyperactivation of the Ras/PKA pathway prevented growth attenuation through ribosome downregulation. We measured the cell volumes of our *cdc-ts AID-BCY1* strains, which we then compared to the cell volumes of *cdc-ts* mutants without the *AID-BCY1* construct. While the mean cell volume of *cdc28-13* cells plateaued, *cdc28-13 AID-BCY1* cells continued to increase in cell volume over the 6 hours (*Figure 5A*, red and dark red, respectively). As before, *cdc20-1* cells were larger than *WT* cells initially, but *cdc20-1 AID-BCY1* cells had an initial mean cell volume twice as large as that of *cdc20-1* and were approximately 600 fL after 6 hours of cell cycle arrest (*Figure 5B*, purple and dark purple, respectively). Cells containing the *cdc15-2* mutant had a mean cell volume of 400 fL after 6 hours of being arrested in the cell cycle (*Figure 5C*, blue). In comparison, *cdc15-2 AID-BCY1* strains grew to be 600 fL after a 6-hour cell cycle arrest, continuing to increase their size exponentially rather than plateauing (*Figure 5C*, dark blue). It is worth noting that cycling *WT AID-BCY1* cells were larger than *WT* cycling (*Figure 5A-C*, black and gray, respectively) upon auxin addition, suggesting that the increase in size may not be specific to *cdc-ts* mutants. All three *cdc-ts* strains with a hyperactive Ras/PKA pathway could no longer regulate their cell size during cell cycle arrests.

**Figure 5.**
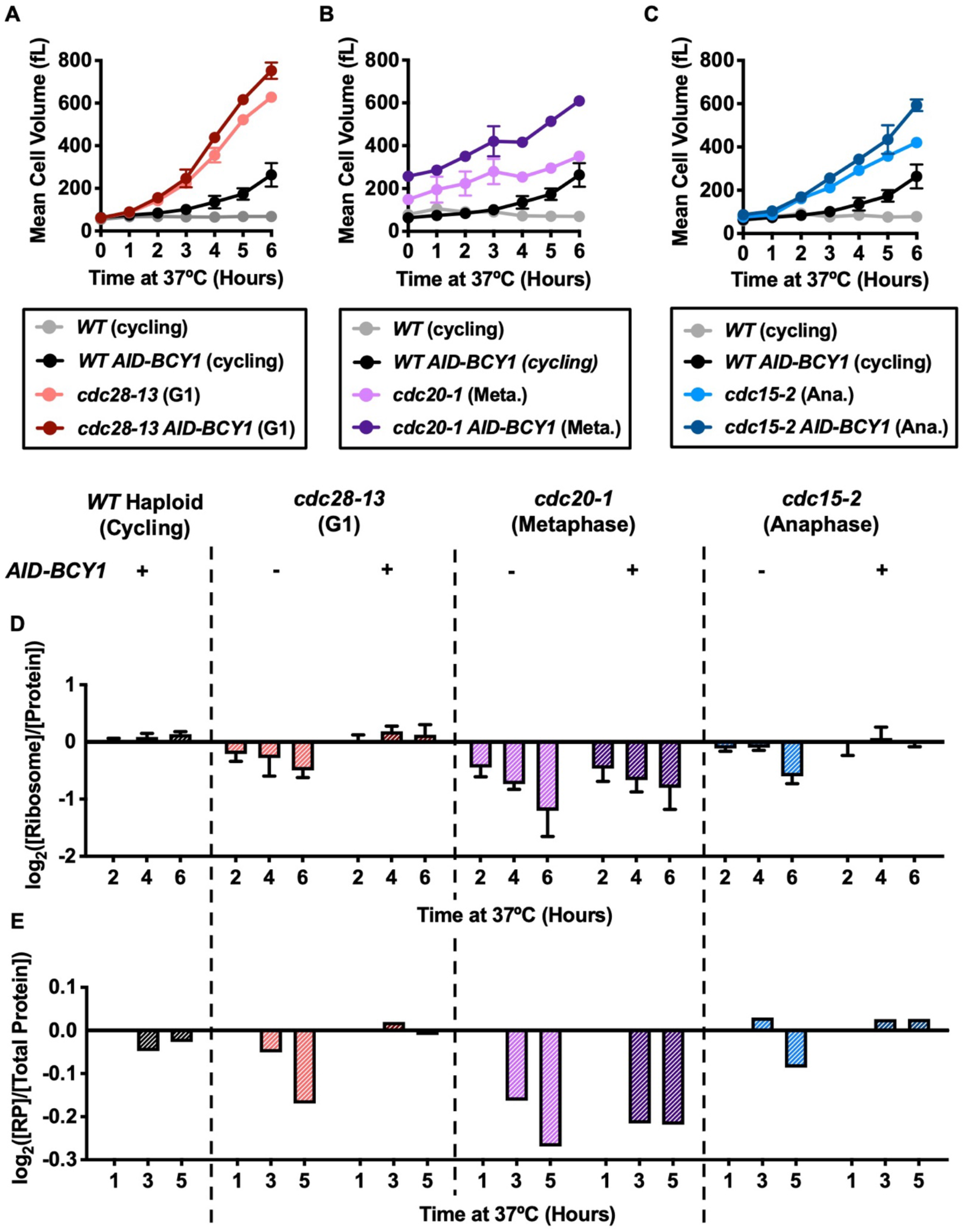
Hyperactivation of the Ras/PKA pathway attenuates ribosome depletion during cell cycle arrest. *WT* (gray, A2587), *WT AID-BCY1* (black, A40439), *cdc28-13* (red, A39000), *cdc28-13 AID-BCY1* (dark red, A40444), *cdc20-1* (purple, A937), *cdc20-1 AID-BCY1* (dark purple, A40499), *cdc15-2* (blue, A2596), and *cdc15-2 AID-BCY1* (dark blue, A40501) cells were grown to log phase in YEPD (supplemented with 138 μL glacial acetic acid for each 1 L YEPD) at 25°C and then shifted to 37°C for 9 hours. *WT* and *WT AID-BCY1* cultures were kept in log phase, termed cycling, at OD_600nm_ 0.2-0.8, by diluting with pre-warmed (37°C) YEPD (supplemented with 138 μL glacial acetic acid for each 1 L YEPD). 500 μM indole-3-acetic acid was added after 2 hours and 4 hours at 37°C. ***(A-C)*** Mean cell volume (fL) was measured for *WT* (gray), *WT AID-BCY1* (black) and ***(A)*** *cdc28-13* (red) and *cdc28-13 AID-BCY1* (dark red), ***(B)*** *cdc20-1* (purple) and *cdc20-1 AID-BCY1* (dark purple), and ***(C)*** *cdc15-2* (blue) and *cdc15-2 AID-BCY1* (dark blue). Error bars represent standard deviation from the mean of experimental replicates. ***(D)*** Protein and ribosome concentrations were quantified using the method described in Terhorst et al. (2020; 17). [Ribosome]/[Protein] was determined. Values were normalized to those of the *WT* cycling samples at each time point and subsequently log_2_ transformed. Error bars represent standard deviation from the mean of experimental replicates. ***(E)*** TMT Proteomics was performed. The fraction of ribosomal proteins (RP) in total protein extracts ([RP]/[Total Protein]) was calculated. Values were normalized to the 1-hour time point in each experiment and subsequently log_2_ transformed.

We next determined whether *cdc-ts* mutants with hyperactive Ras/PKA were able to selectively downregulate ribosom fraction of the proteome, through decreased [Ribosome]/[Protein] levels. *cdc28-13* exhibited a selective downregulation of ribosomes (*Figure 5D*, red), but *cdc28-13 AID-BCY1* cells no longer decreased [Ribosome]/[Protein] (*Figure 5D*, dark red). While ribosome downregulation appeared to occur in *cdc20-1 AID-BCY1* cells, there was more [Ribosome]/[Protein] in *cdc20-1 AID-BCY1* cells than in *cdc20-1* at each time point (*Figure 5D*, dark purple and purple, respectively). As expected, *cdc15-2* cells decreased [Ribosome]/[Protein] over the 6-hour cell cycle arrest (*Figure 5D*, blue), but *cdc15-2 AID-BCY1* cells no longer change [Ribosome]/[Protein] when arrested (*Figure 5D*, dark blue). Importantly, there was also no change in [Ribosome]/[Protein] for *WT AID-BCY1* cells in comparison to *WT* cells, suggesting that hyperactivation of the Ras/PKA pathway does not cause an increase in the ribosomal fraction of the proteome in every strain we studied but instead uniquely in *cdc-ts* arrested cells (*Figure 5D*, black). As before, we verified the changes in the ribosomal fraction of the proteome by measuring [RP]/[Total Protein] with TMT Proteomics (*Figure 5E*). We conclude that while preventing ESR activation, *cdc-ts* mutants with hyperactive Ras/PKA pathways can no longer attenuate their size through specific ribosome downregulation.

## Discussion

Previous work from our lab had thoroughly investigated the change in cell volume of *cdc-ts* mutants that were arrested in the cell cycle, but the mechanism behind cell volume plateauing in those mutant strains remained unknown (9). Here we confirmed that after prolonged cell cycle arrest, cells of three independent *cdc-ts* strains were able to regulate their cell volume and halt growth. Because ribosome synthesis has been shown to regulate cell size, we studied the protein and ribosome content of the *cdc-ts* cells using two methods (4, 10). There was a selective ribosome downregulation in each strain, independent of the stage of the cell cycle arrest. Ribosome downregulation was observed upon cell cycle arrest using three different methods that mimicked naturally occurring cell cycle arrests: alpha factor addition, entrance into stationary phase, and TORC1 inhibition by rapamycin. Interestingly, while alpha factor addition and entrance into stationary phase caused cells to decrease their size, rapamycin addition led to increased cell volume in both *WT* haploid and diploid strains. Each cell cycle arrest studied activated the ESR, a transcriptional fingerprint known to downregulate ribosomal proteins and biogenesis factors. We also observed that cells exhibited a greater induction of the ESR when they were unable to make their own amino acids. When we prevented ESR activation in *cdc-ts* cells by hyperactivating the Ras/PKA pathway we observed decreased cell viability. Additionally, cells with a hyperactive Ras/PKA pathway were unable to attenuate their size through ribosomes downregulation. Taken together, we conclude that ESR activation is required for *cdc-ts* cells in order to regulate their volume and survive cell cycle arrests through downregulation of ribosome content.

Our findings uncover new questions about how *cdc-ts* mutants regulate their cell volume when arrested. The cause of ESR activation in *cdc-ts* arrests remains unknown but poses a complex question to answer because each strain studied may activate the ESR through different mechanisms, which might be related to the stage in which the cells are arrested, or the mutation itself. Therefore, it is important to continue to study multiple *cdc-ts* strains in comparison to other cell cycle arrests, as we have done here. In our studies, the ESR activation in naturally occurring cell cycle arrests, alpha factor and rapamycin addition and entrance into stationary phase, suggested three possible mechanisms of ESR activation in *cdc-ts* cells: the TORC1, MAPK, and/or Ras/PKA pathways. Inhibiting the TORC1 pathway through rapamycin addition caused cells to increase in size, likely due to vacuole growth (28). Goranov et al. (2013, 23) hyperactivated the TORC1 pathway in *cdc28-4* cells, a temperature-sensitive allele of *CDC28*, and the TORC1 hyperactivation appeared to cause cells to be even smaller than size attenuated *cdc28-4* cells during the arrest. We confirmed this finding in *cdc28-13* cells as well (data not shown). Taken together, those results suggest that TORC1 activity is antagonistic to size attenuation in *cdc-ts* arrests. Another group of pathways that may be involved in cell size regulation in *cdc-ts* mutants is the MAPK pathways. ESR activation through alpha factor addition suggests that MAPK pathways could be activated or inactivated in response to *cdc-ts* arrests. Specifically, the cell wall integrity pathway may be activated in *cdc-ts* arrests, particularly *cdc28-13*, since cell volume increases rapidly throughout the arrest, causing significant stress within the cell (36). Another likely candidate is Ras/PKA pathway inactivation since its hyperactivation prevented ESR activation and size attenuation through ribosomal downregulation. These pathways, and many others, are highly interconnected, so it would not be surprising if multiple pathways are involved (37).

We suspect that ribosomes may be disassembled, if not wholly degraded. When we measured the ribosomal fraction of the proteome, we used two independent methods: one involving purifying assembled ribosomes and the other involving measuring ribosomal proteins of whole cell lysates. When we measured the rRNA concentration of purified assembled ribosomes in the ribosome quantification method, there was a more dramatic decrease in the ribosomal fraction of the proteome than when we measured ribosomal proteins in whole cell lysates (*Figure 1B-C*). As previously mentioned, this discrepancy could be simply be due to one of the many other differences between these two methods, but an alternative, more interesting explanation is that ribosomes may be disassembled when *cdc-ts* mutants are arrested in the cell cycle, and various components may be degraded downstream the disassembly. If disassembly and/or degradation were occurring upon arrest in the cell cycle, ribosomal components may be used in the synthesis of other macromolecules needed for survival as we know that ESR activation leads to increased biosynthesis of certain carbohydrates, fatty acids, and the cell wall (12). In this case, the cell’s ribosomes would be used as supply key elements needed to survive cell cycle arrests.

Finally, questions remain concerning the mechanism by which cells continue to be viable after a prolonged arrest in the cell cycle. The differences in cell viability between *cdc-ts* mutants with and without a hyperactive Ras/PKA pathway were striking. When cells were no longer able to activate the ESR, cell viability decreased dramatically (*Figure 4G-H*). We did not know which components of the ESR were responsible for cell viability or what their respective contributions to survival were. Specifically, we remain curious about the contribution of size attenuation and ribosome downregulation to cell survival in *cdc-ts* arrests. By preventing cells from growing too large and diluting their cytoplasm, ribosomes may be involved in a key mechanism in preventing cell death. Using *cdc-ts* cells as a model system, we have shown that the ESR and ribosome downregulation have profound effects on cell physiology and survival.

## Acknowledgements

We thank Frank Solomon, Monica Boselli, Alexi Goranov, and Summer Morrill for comments, the MIT BioMicroCenter for RNA-Seq, the MIT Swanson Biotechnology Center for proteomics, and Amy Ikui for exciting discussions about our results. This work was supported by NIH grants CA206157 and GM118066 to A.A., who was an investigator of the Howard Hughes Medical Institute, the Paul F. Glenn Center for Biology of Aging Research at MIT, and the Ludwig Center at MIT.

## Data Deposition

RNA-Seq data has been deposited in the Gene Expression Omnibus (GEO) database (accession no. TBD).

## Materials and Methods

### Yeast strains and growth conditions

Yeast strains used in this study are of the W303 background and described in *Table S*1. Unless otherwise noted, strains were grown in YEPD complete media. *AID-BCY1* strains were grown in YEPD supplemented with 138 μL glacial acetic acid for each 1 L YEPD. 500 μM indole-3-acetic acid (Sigma-Aldrich) was used as auxin in the *AID-BCY1* experiments.

### Mean Cell Volume Measurements

1 mL samples of culture were diluted 1:100 with Isoton II Diluent (Beckman Coulter) and measured on a Beckman Multisizer 3 Coulter Counter to produce a histogram of the population’s cell volumes. Values above the half-maximal cell count were used to calculate the mean cell volume of the culture.

### Protein and Ribosome Quantification

Protein and ribosome quantification were performed as in Terhorst et al. (2020, 17).

### TMT Proteomics

TMT Proteomics was performed by the method described in Rose et al. (2016, 40).

### GEM Diffusion Measurements

GEM diffusion measurements were performed as in Delarue et al. (2018; 21).

### RNASeq

RNASeq was performed as in Terhorst et al. (2020, 17). All sequencing was done using an Illumina HiSeq2000.

### Data Processing and Single-Sample Gene Set Enrichment Analysis

Data processing and ssGSEA was performed as in Terhorst et al. (2020, 17).

### Cell Viability Measurements

1 mL samples of culture were diluted 1:100 diluted with Isoton II Diluent (Beckman Coulter) and used to determine cell concentration of the culture using a Beckman Multisizer 3 Coulter Counter. 300 cells were plated on YEPD agar plates and grown at 25°C until colonies appeared, approximately 3-5 days. Colonies were counted by hand, and values were normalized to the 2-hour time point of each experiment.

### Data Availability

The data discussed in the paper are included in the figures and SI Appendix. RNA-Seq data has been deposited in the Gene Expression Omnibus (GEO) database (accession no. TBD).

