## Supplementary Data for "The environmental stress response regulates ribosome content in cell cycle-arrested *S. cerevisiae*"

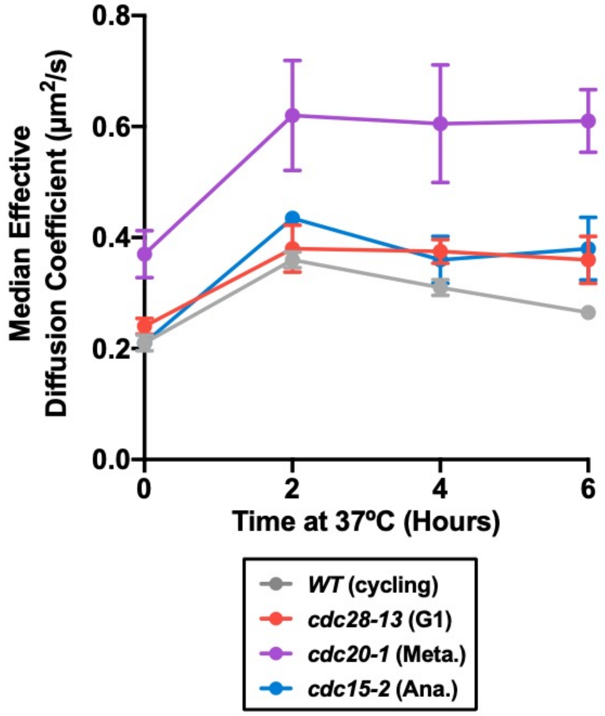


**Figure S1. Macromolecular crowding in *cdc-ts* mutants arrested in the cell cycle.**
*WT* (gray, A2587), *cdc28-13* (red, A39000), *cdc20-1* (purple, A937), and *cdc15-2* (blue, A2596) cells were grown to log phase in YEPD at 25ºC and then shifted to 37ºC for 6 hours. *WT* cultures were kept in log phase, termed cycling, at $OD_{600nm}$ = 0.2-0.8, by diluting with pre-warmed (37ºC) YEPD. GEM diffusion was performed as in Delarue et al. (2018; 21) to calculate median diffusion coefficients. Error bars represent standard deviation from the mean of experimental replicates.

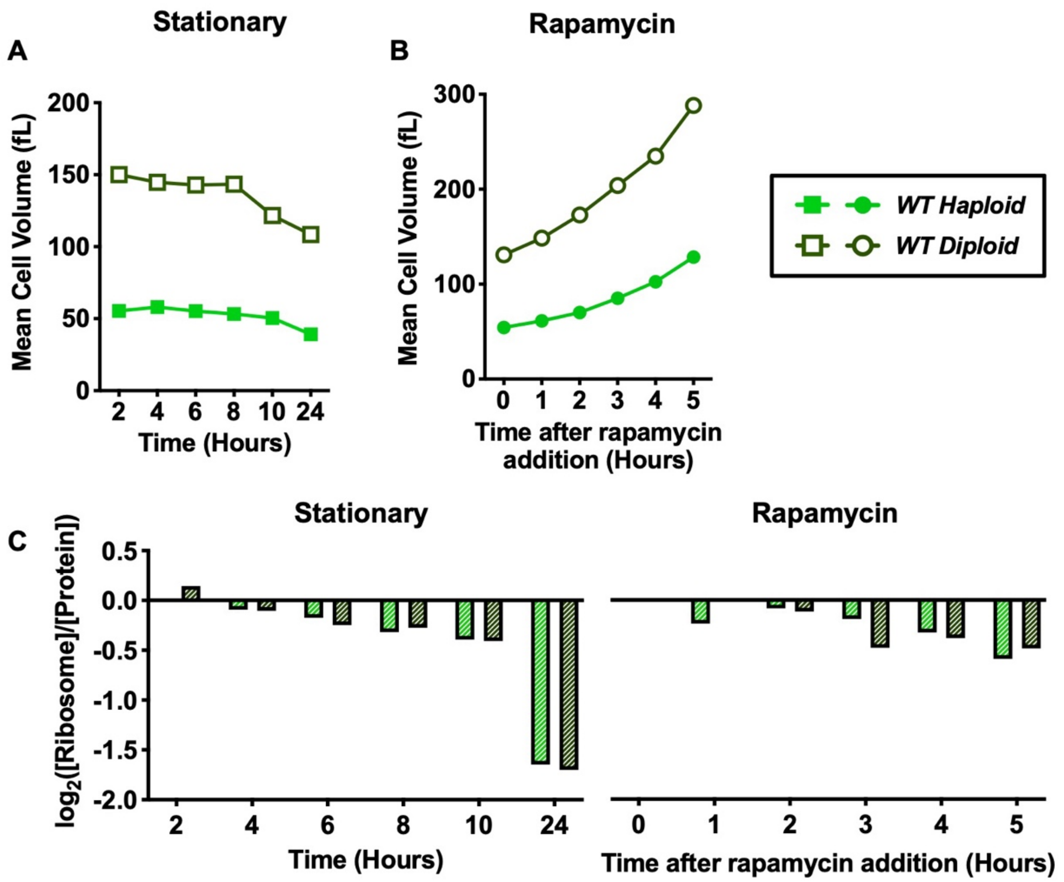


**Figure S2. Comparison of protein and ribosome quantification in haploid and diploid *WT* cells.**
For stationary phase experiments, *WT* haploid (green, A2587) and *WT* diploid (dark green, A33728) cells were grown in YEPD for 24 hours at 30ºC. For rapamycin experiments, *WT* haploid (green, A2587) and *WT* diploid (dark green, A33728) cells were grown to log phase in YEPD at 30ºC. 5 nM rapamycin was added to *WT* haploid cells, and 2.5 nM rapamycin was added to *WT* diploid cells. Cells were grown for 5 hours at 30ºC in the presence of rapamycin. ***(A-B)*** Mean cell volume (fL) was measured for *WT* haploid (green) and *WT* diploid (dark green) cells grown ***(A)*** into stationary phase (squares) and ***(B)*** in the presence of rapamycin (circles). ***(C)*** Protein and ribosome concentrations were quantified using the method described in Terhorst et al. (2020; 17). [Ribosome]/[Protein] was determined. Values were normalized to the 2-hour time point in stationary experiments and to the 0-hour time point in rapamycin experiments and subsequently $\log_2$ transformed.

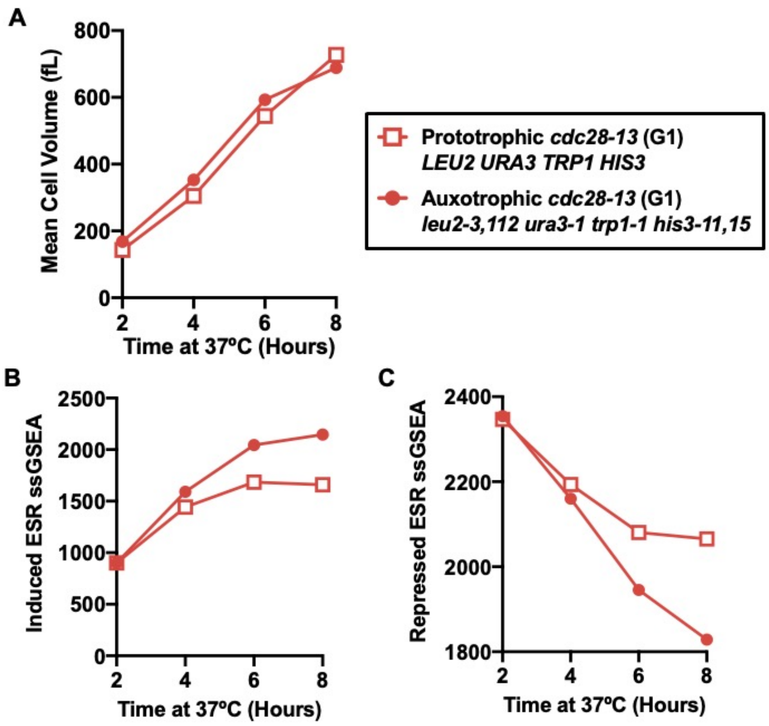


**Figure S3. Comparison of ESR activation in prototrophic *cdc28-13* cells and auxotrophic *cdc28-13* cells.**

Prototrophic *cdc28-13* (red squares, A41270) and auxotrophic *cdc28-13* (red circles, A17896) were grown to log phase in YEPD at 25ºC. Cells were synchronized with alpha factor then shifted to 37ºC for 6 hours.

***(A)*** Mean cell volume (fL) was measured for prototrophic *cdc28-13* cells (red squares) and auxotrophic *cdc28-13* cells (red circles).

***(B-C)*** RNA-Seq samples were collected, and gene expression data were analyzed by calculating ssGSEA projection values for the (***B***) induced ESR and ***(C)*** repressed ESR.

| Name | Number | Genotype | Source |
|---|---|---|---|
| *WT* Haploid | A2587 | *MATa, ade2-1, leu2-3, ura3, trp1-1, his3-11,15, can1-100, GAL, psi+* | Nasmyth Lab |
| *cdc28-13* | A39000 | *MATa, ade2-1, leu2-3, ura3, trp1-1, his3-11,15, can1-100, GAL, psi+, cdc28-13::URA3* | Amon Lab |
| *cdc20-1* | A937 | *MATalpha, cdc20-1, ura3, trp1, leu2, his3, ade2, can1* | Nasmyth Lab |
| *cdc15-2* | A2596 | *MATa, cdc15-2, leu2-3, ura3, trp1-1, omns, ade1* | Nasmyth Lab |
| *WT GEM* | LH4415 | *MATa, ade2-1, leu2-3, ura3, trp1-1, his3-11,15, can1-100, GAL, psi+ HIS3::pINO4-PfV-Sapphire* | Holt Lab |
| *cdc28-13 GEM* | LH4416 | *MATa, ade2-1, leu2-3, ura3, trp1-1, his3-11,15, can1-100, GAL, psi+, cdc28-13::URA3 HIS3::pINO4-PfV-Sapphire* | Holt Lab |
| *cdc20-1 GEM* | LH4417 | *MATalpha, cdc20-1, ura3, trp1, leu2, his3, ade2, can1 HIS3::pINO4-PfV-Sapphire* | Holt Lab |
| *cdc15-2 GEM* | LH4418 | *MATa, cdc15-2, leu2-3, ura3, trp1-1, omns, ade1 HIS3::pINO4-PfV-Sapphire* | Holt Lab |
| *bar1Δ* | A2589 | *MATa, bar1::HisG, leu2-3, ura3, trp1-1, his3, ade2, can1-100, GAL, psi+,* | Nasmyth Lab |
| *WT* Diploid | A33728 | *MATa/alpha, ade2-1, leu2-3, ura3, trp1-1, his3-11,15, can1-100, GAL, psi+* | Fink Lab |
| *cdc28-13* Auxotroph | A17896 | *MATa, cdc28-13, ADE2, leu2-3, ura3, trp1-1, his3-11,15, can1-100, GAL, psi+* | Amon Lab |
| *cdc28-13* Prototroph | A41270 | *MATa, cdc28-13, ADE2, URA3, TRP1, HIS3, can1-100, GAL, psi+, cdc28-13, leu2::4xSTRE-GFP:LEU2* | Amon Lab |
| *WT AID-BCY1* | A40439 | *MATa, ade2-1, leu2-3, ura3, trp1-1, his3-11,15, can1-100, GAL, psi+, KanMX:pRFA1:9Myc-AID-BCY1, leu2::pTEF1-osTIR::LEU2* | Amon Lab |
| *cdc28-13 AID-BCY1* | A40444 | *MATa, ade2-1, leu2-3, ura3, trp1-1, his3-11,15, can1-100, GAL, psi+, cdc28-13::URA, KanMX:pRFA1:9Myc-AID-BCY1, leu2::pTEF1-osTIR::LEU2* | Amon Lab |
| *cdc20-1 AID-BCY1* | A40499 | *MATa, ade2-1, leu2-3, ura3, trp1-1, his3-11,15, can1-100, GAL, psi+, cdc20-1, KanMX:pRFA1:9Myc-AID-BCY1, leu2::pTEF1-osTIR::LEU2* | Amon Lab |
| *cdc15-2 AID-BCY1* | A40501 | *MATa, ade2-1, leu2-3, ura3, trp1-1, his3-11,15, can1-100, GAL, psi+, cdc15-2, KanMX:pRFA1:9Myc-AID-BCY1, leu2::pTEF1-osTIR::LEU2* | Amon Lab |

**Table S1. Yeast strains used in this study.** Description of the strain names, numbers, genotypes, and source used in this paper.
